# Effects of culture medium and temperature on fungal growth of *Mycena* and marasmioid isolates and *in vitro* symbiotic culture in mycoheterotrophic orchid, *Gastrodia pubilabiata*

**DOI:** 10.1101/2021.10.07.463587

**Authors:** Akane Shimazaki, Kana Higaki, Kento Rammitsu, Yuki Ogura-Tsujita

**Author notes:** Corresponding author, E-mail address (Y. Ogura-Tsujita).

## Abstract

In vitro symbiotic culture of *Gastrodia pubilabiata* seeds were conducted with the fungal isolates from *G. pubilabiata* roots. We obtained five fungal isolates, which belonged to *Mycena*, Marasmiaceae and Omphalotaceae. Firstly, optimal temperature and culture medium for subculture of these fungal isolates were examined. All five isolates grew the fastest on malt extract agar medium. Mycelial growth rate was highest at 25 °C between 10 °C and 40 °C. Secondly, we evaluated suitable culture vessels and organic materials for symbiotic culture. Seeds germinated well in petri dishes with *Quercus* leaf disc on water agar medium, and the seed germination process was well observed without dense mycelium. The most developed seedlings were found in glass bottles filled with Japanese cedar leaves, but densely grew mycelium prevent accurate seedling counts. Leaves of *Quercus*, Japanese cedar and bamboo were used as organic materials for symbiotic culture. All three leaves induced seed germination with *Mycena* and Marasmiaceae fungi, but material types affected subsequent seedling growth. Our method will contribute to understanding the mycorrhizal association of *Gastrodia* species and also other mycoheterotrophic plants.

## Introduction

Mycoheterotrophic plants, which are leafless and achlorophyllous, depend entirely on their mycorrhizal fungi for their carbon supply (Leake 1994). Symbiotic relationship between mycoheterotrophic plants and their mycorrhizal fungi are still not well understood. Despite many studies have identified the fungal taxa of mycorrhizal fungi associating with mycoheterotrophic plants (Waterman et al. 2013), the physiological mechanisms underlying plant and fungus interactions in mycoheterotrophic association, such as nutrient transport and recognition between plant and fungus, are still unclear. In vitro symbiotic culture system with plant and fungus will contribute to understanding physiology of mycoheterotrophy. Furthermore, in vitro symbiotic culture has been used as a method to evaluate symbiotic compatibility between plant and mycorrhizal fungus (Rasmussen, 2002). Specificity for mycorrhizal fungus is highly diverge across plant phylogeny (Wang et al. 2021), which is interesting to understand the evolution of mycoheterotrophic plants. In ecological aspect, a life cycle of mycoheterotrophic plants is less understood because these plants spent most of their life cycle underground and appear aboveground only in flowering season. In vitro culture will reveal an unknown life cycle of mycoheterotrophic plants. However, the study of in vitro symbiotic culture in mycoheterotrophic plants is quite limited.

Mycoheterotrophic plants that associate with ectomycorrhizal or arbuscular mycorrhizal fungi are difficult to cultivate artificially, because these fungi are obligately biotrophic and depend on autotrophic plants for their carbon supply. Thus, chlorophyllous host plant is required to maintain ectomycorrhizal and arbuscular mycorrhizal fungi, which causes difficulty in artificial cultivation. On the other hand, mycoheterotrophic plants that associate with free-living saprotrophic fungi, have been cultivated in several plant species, such as *Didymoplexis micradenia* (former *D. minor*; Burgeff 1932) and *Epipogium roseum* (Yagame et al. 2007).

Fully mycoheterotrophic *Gastrodia* is one of the most divergent mycoheterotrophic genus and mainly associated with litter- or wood-decaying fungi (Ogura-Tsujita et al. 2021). Symbiotic culture of *Gastrodia* species has been achieved by in vitro system or field cultivation, mainly in medicinally important crop, *G. elata* (Xu and Guo 2000). Since *G. elata* is large in size and difficult to complete its life cycle in vitro, smaller species is preferable for in vitro culture experiments. However, such culture techniques have been less applied to other *Gastrodia* species. Successful in vitro seed germination of *G. verrucosa* with an unidentified clamp bearing fungus (Tashima et al. 1978) suggests that in vitro symbiotic culture technique can be applied to other *Gastoridia* species. *G. pubilabiata* has been successfully cultured in various studies and flowering was achieved under cultivation (Shimaoka et al. 2017). The seeds of *G. pubilabiata* were germinated in a plastic box containing leaf litter or log collected from its habitats by using naturally inhabiting fungi within the organic materials (Higaki et al. 2017, Shimaoka et al. 2017). Furthermore, in vitro symbiotic germination was succeeded in *G. pubilabiata* by using a fungal isolate from *G. confusa* (Umata et al. 2000) and also in *G. nipponica* by using the isolates from fungal fruit bodies (Umata and Nishi 2010). However, these isolates were unidentified in both studies, and the culture conditions for maintaining fungal isolates, such as culture media and temperature, were not examined. Furthermore, the culture vessels used in the previous studies varied among studies, such as petri dish (Park et al. 2012), glass bottle (Higaki et al. 2017) and three-compartment petri dish (Umata and Nishi 2010), and the suitable culture vessel has yet to be examined. As the mycorrhizal fungi of *Gastrodia* species are mainly leaf litter- or wood-decaying fungi, organic materials, such as fallen leaves and wood logs, have been used for symbiotic culture as a substrate for fungal isolates (Ogura-Tsujita et al. 2021). Pieces of dead *Quercus* leaves has been used in the symbiotic culture of *G. elata* seeds with *Mycena* fungi (Park et al. 2012). Bamboo (Tashima et al. 1978) and Japanese cedar, *Cryptomeria japonica*, materials (Higaki et al. 2017, Shimaoka et al. 2017) were also used for *Gastrodia* culture. Since the suitable organic material may vary among fungal species, examination of appropriate organic material for used fungal isolates is required for symbiotic culture.

To establish an in vitro culture system in *G. pubilabiata*, we attempt to isolate and identify the mycorrhizal fungi of this species and to culture the *G. pubilabiata* seeds with these isolates. Firstly, we evaluated the suitable culture medium and temperature for subculturing fungal isolates. Secondly, we evaluated a suitable culture vessel for in vitro symbiotic culture of *G. pubilabiata* among three vessels, petri dish, glass bottle and three-compartment petri dish. We also examined three organic materials, the leaves of *Quercus*,bamboo and Japanese cheder, for symbiotic culture, because *G. pubilabiata* is mainly associated with leaf litter-decaying fungi and this orchid often found in evergreen broadleaved forest, bamboo thicket and Japanese cheder forest (Kinoshita et al. 2016).

## Materials and Methods

### Fungal isolation

Malt extract agar medium (malt extract 20g, agar 15g, water 1L) supplemented with 50 mg/L streptomycin and 50 mg/L tetracycline was used to isolate the mycorrhizal fungi of *Gastrodia pubilabiata*. Mycorrhizal roots were collected from adult plants in natural habitat or seedlings cultivated by using leaf litter obtained from natural habitats (Higaki et al. 2017). Root tips were washed with sterilized water and clashed within distilled water. Released fungal coils were collected by a micropipet under stereomicroscope and dropped on the isolation medium. The plates were incubated at 25 °C under dark condition for three to seven days. Elongated fungal hyphae were transferred onto new 2% malt extract agar medium to obtain pure cultures.

### Molecular identification of fungal isolates

Obtained isolates were grouped by colony morphology and representative isolates of each group were identified. Fungal DNA was extracted from the mycelial pieces according to Izumitsu et al. (2012). PCR amplification and sanger sequencing were carried out as described by Ogura-Tsujita and Yukawa (2008). Internal transcribed spacer (ITS) sequences were amplified using the primer pair ITS1F/ITS4 (White et al. 1990; Gardes and Bruns 1993). The obtained sequences were analyzed using a BLAST search against the GenBank sequence database to find the closest matching sequence. The representative ITS sequences of each fungal group were deposited in the DNA Data Bank of Japan (DDBJ; see Table 1). Isolates F68 and F69 were already published in Higaki et al. (2017).

**Table 1.**
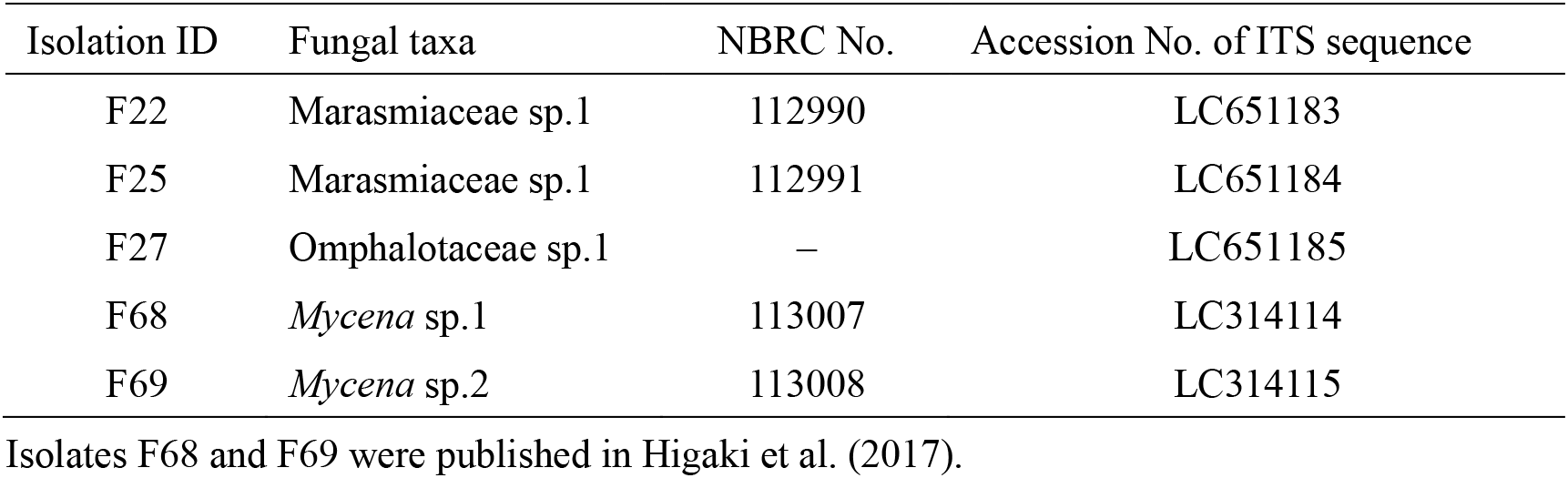
Fungal isolates from *Gastrodia pubilabiata* roots.

### Fungal subculture medium

Five types of culture media were evaluated for the subculture of fungal isolates; 1) potato dextrose agar (PDA) medium, 2) 2% malt extract agar medium, 3) modified Melin–Norkrans medium (MMN; sucrose 10.0 g, NaCl 0,025g, 1.0 g, KH_2_PO_4_ 0.5 g, (NH_4_)_2_HPO_4_ 0.25g, MgSO_4_·7H_2_O 0.15 g, CaCl_2_·2H_2_O 0.05 g, FeCl_3_ 1.2 mg, thiamine hydrochloride 0.1 mg, malt extract 3.0 g, yeast extract 2.0 g, agar 15.0 g, distilled water 1,000 ml; Marx 1969), 4) Murashige and Skoog (MS) medium (Duchefa Biochemie BV), 5) 1.5% agar medium. The former three media are standard media for fungal culture, and 1.5% agar medium is a negative control without nutrient. A MS medium is a standard medium for plant tissue culture and examined fungal growth to confirm whether this medium is appropriate for symbiotic culture. Five fungal isolates (F22, F25, F27, F68, F69; Table 1) were used for the examination. Subcultured hyphal tips were transferred onto 90 mm petri dishes containing 20 mL each medium and cultured at 25 °C under dark condition. Four replicates were used for each medium type. The diameter of the fungal colonies was recorded for maximum 27 days.

### Temperature for fungal subculture

Three isolates (F25, F27, F68) were used. Subcultured hyphal tips were transferred onto 2% malt extract agar medium and cultured at 10, 20, 25, 30 and 40 °C under dark condition. Four replicates were used for each temperature. Colony diameter was measured every two to three days and fungal growth rate (mm/day) were calculated.

### Culture vessels for symbiotic culture

Seeds of *G. pubilabiata* were collected from the matured fruits in the habitat and dried with silica gel in the glass bottle. Dried seeds were stored at 5 °C until use. The seeds were sterilized in 5% Ca(ClO)2 with 200 μL tween20 for 15 minutes and rinsed three times with sterilized distilled water, and spread on the medium. Maximum four fungal isolates (F22, F25, F68, F69; Table 1) were used for the examination. Fungal isolates were subcultured on 2% malt extract agar medium two to three weeks before symbiotic culture. We evaluated three types of culture vessels; 1) a petri dish with *Quercus* leaf disc on 1.5% water agar medium (Fig. 1A), 2) a glass bottle containing *Cryptomeria japonica* leaves (Fig. 1B), 3) a three-compartment petri dish containing PDA medium and 1.5% water agar medium (Fig. 1C). Dead leaves of *Quercus glauca* and *Cryptomeria japonica* were autoclaved at 121°C for two hours, dried for three days at 70 °C and kept at room temperature until use. In the first method, we followed the technique by Park et al. (2012) which induced in vitro seed germination of *G. elata* with *Mycena* fungi. The sterilized *Quercus* leaves were cut into 2 × 2 cm discs (0.03g) and autoclaved again at 121°C for 15 minutes before use. Sterilized discs were immersed in sterile water for five minutes and placed on the subcultured fungal plates to colonize the discs with fungal isolates. After 15 days culture, a disc was placed on the center of the 90 mm petri dish containing 20 mL 1.5% agar medium (Fig. 1A). About 50-100 seeds were sown around leaf disc. For the second method, we followed the methods by Higaki et al. (2017) which successfully induced *G. pubilabiata* seed germination by using *Cryptomeria* leaves collected from natural habitat. The 450 mL glass bottle with pumice on the bottom were filled with sterilized *Cryptomeria* leaves (Fig. 1B) and autoclaved at 121°C for 15 minutes. To moisten the leaves, 30 mL sterilized water were added to the bottle. Hyphal tips were subcultured in the 100 mL flask containing 2% malt extract liquid media at 100 rpm. After 10 days of subculture, the fungal colonies were ground in a sterile mortar and 10 mL suspension was added to the bottle. About 50-200 seeds were sown on each bottle. The third method followed Umata and Nishi (2010) which induced in vitro seed germination of *G. nipponica* with a fungal isolate from fungal fruit body. We used a 100 mm three-compartment petri dish with PDA medium in one compartment and 1.5% water agar medium in other two compartments (Fig. 1C). Fungal inoculum was placed on PDA medium and *G. pubilabiata* seeds were spread on 1.5% water agar media. All vessels were cultured at 25°C in the dark. About 50 seeds were sown on each plate. Four replicates were made for each vessel. After two months of culture, the number of germinated seeds was counted in five stages as shown in Fig. 2 (Stage 0, no germination; Stage 1, enlarged protocorm <0.2 mm in length; Stage 2, elongated seedling between 3 mm and 1 cm in length; Stage 3, elongated seedlings over 1 cm; Stage 4, tuber development).

**Fig. 1.**
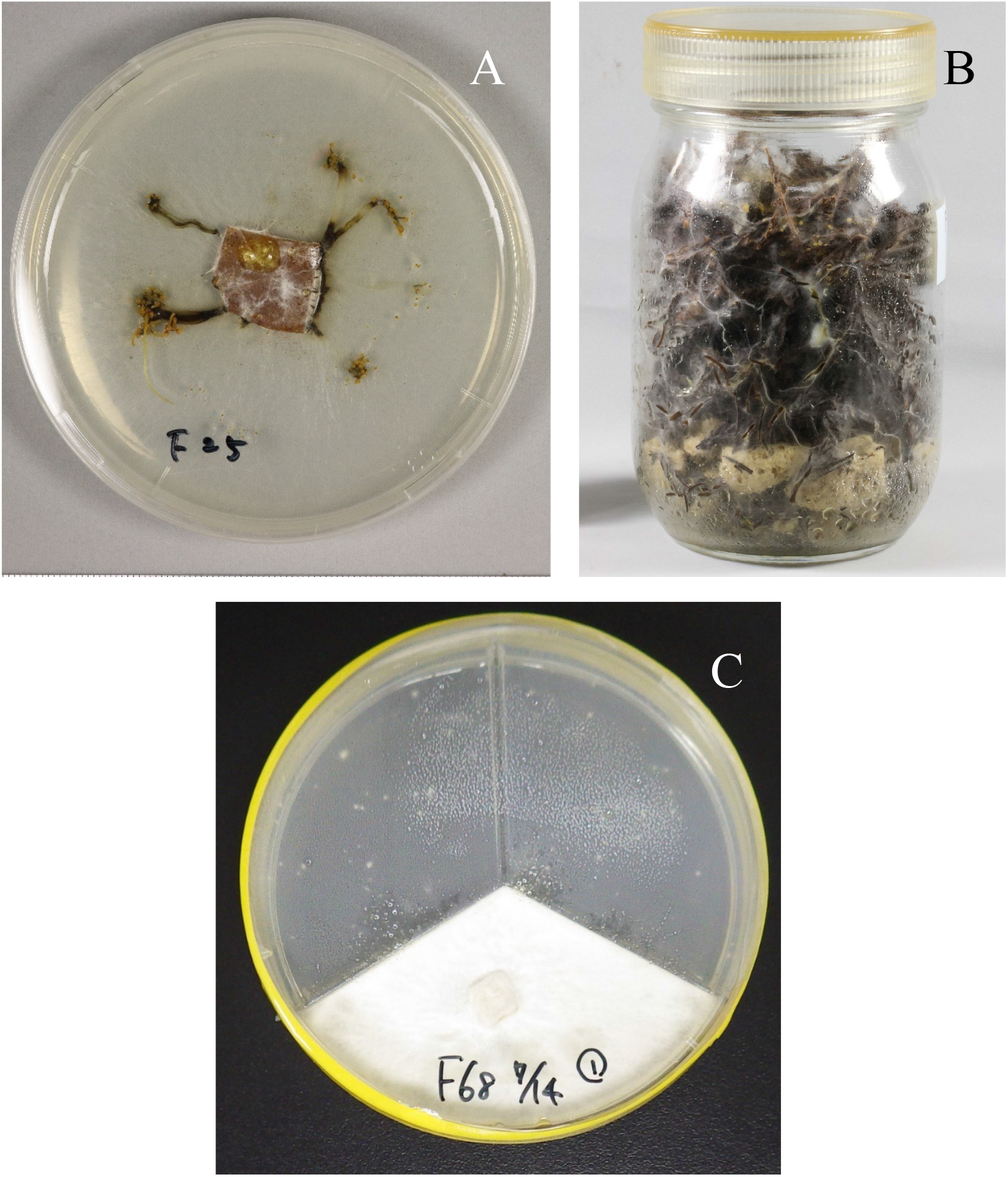
Symbiotic culture of *Gastrodia pubilabiata*. Three different culture vessels used in this study; A: petri dish with *Quercus* leaf disc, B: glass bottle with *Cryptomeria* leaves, C: three-compartment petri dish with fungal inoculum on PDA medium in one compartment and seeds on 1.5% water agar medium in other two compartments.

**Fig. 2.**
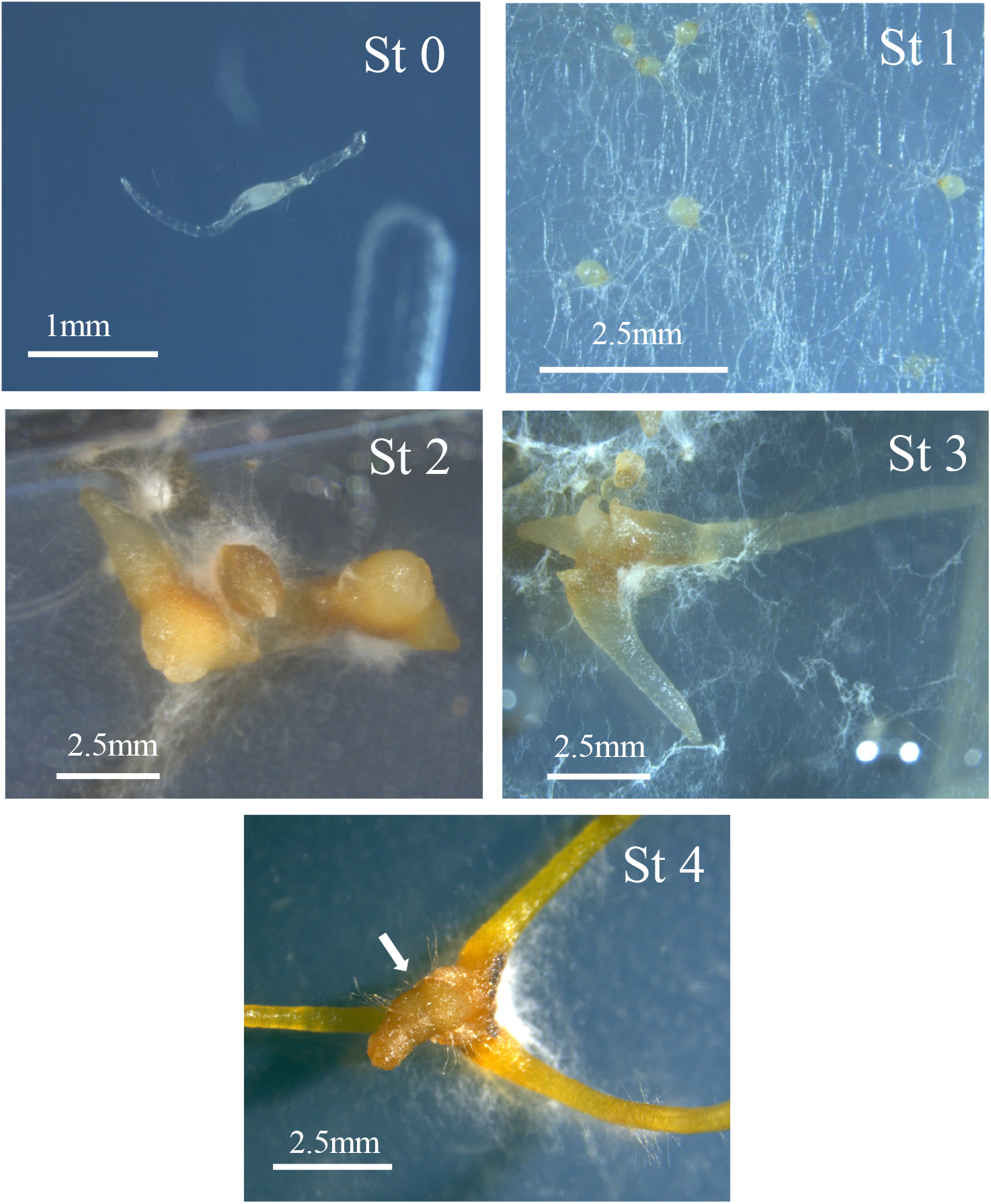
Seed germination stages; St 0: no germination, St 1: elongated protocorm <0.2 mm in length, St 2: elongated seedlings between 3 mm and 1 cm in length, St 3: elongated seedlings over 1 cm, St 4: tuber development (arrow shows tuber).

### Organic materials for symbiotic culture

We used the dead leaves of three plants, *Quercus glauca, Cryptomeria japonica* (Japanese cedar) and *Phyllostachys edulis* (bamboo), for a substrate of fungal isolates. Two fungal isolates (F25 and F68; Table 1) were employed for the experiment. All dead leaves were autoclaved at 121°C for two hours, dried for three days at 70 °C and kept at room temperature until use. The leaves were cut into 0.03g discs and autoclaved again at 121°C for 15 minutes before use. The subsequent procedure was performed in the same protocol as the first methods described above. About 50-200 seeds were sown on each plate and four replicates were made for each material. The number of germinated seeds was counted in five stages after two months of culture.

## Results and Discussion

### Fungal isolation and identification

A total of five fungal strains were isolated from *Gastrodia pubilabiata* roots (Table 1). These strains belonged to Marasmiaceae, Omphalotaceae and *Mycena*. The ITS sequences of F22 and F25 shared 99% identity, and both sequences had 99% similarities with *Marasmiellus* fungi from *Gastrodia nipponica* (LC013366) and *G. pubilabiata* (LC013342) roots (Kinoshita et al. 2016). These *Marasmiellus* fungi were shown to belong to a campanelloids clade within Marasmiaceae in Ogura-Tsujita et al. (2021). The sequence of F27 showed the highest similarity (96%) to that from fungal fruit body of *Marasmius* sp. (LC504843). Furthermore, the isolate F27 had 91% sequence similarity with *Marasmiellus ramealis* (DQ450031), which is included in *Marasmiellus* clade within Omphalotaceae (Oliveria et al. 2019; Ogura-Tsujita et al. 2021). Sequences from F68 and 69 shared 97%identity, and those two sequences shared more than 97% similarities to *Mycena* fungi identified from *G. nipponica* (LC013372) and *G. pubilabiata* (LC013346) roots (Kinoshita et al. 2016).

### Fungal subculture medium

All fungal isolates grew fastest on the malt extract agar medium, suggesting that this medium would be the best for the fungal subculture of *Mycena*,Marasmiaceae and Omphalotaceae isolates (Fig. 3). The fungal growth rate varied among the isolates. The isolate F25 grew fastest and took seven days to reach the edge of the petri dish, while the isolate F69 took 16 days. The growth rate of mycelium differed greatly depending on the medium type. The colony of isolate F68 reached the edge of the petri dish in 12 days on malt extract agar medium, but it took more than 20 days on other media. Faster growth of *Mycena* mycelium on malt extract agar medium than on PDA medium was also reported by Hong et al. (2002). Culture on MS medium strongly inhibits fungal growth in all fungal isolates, probably due to too much nitrogen salts.

**Fig. 3.**
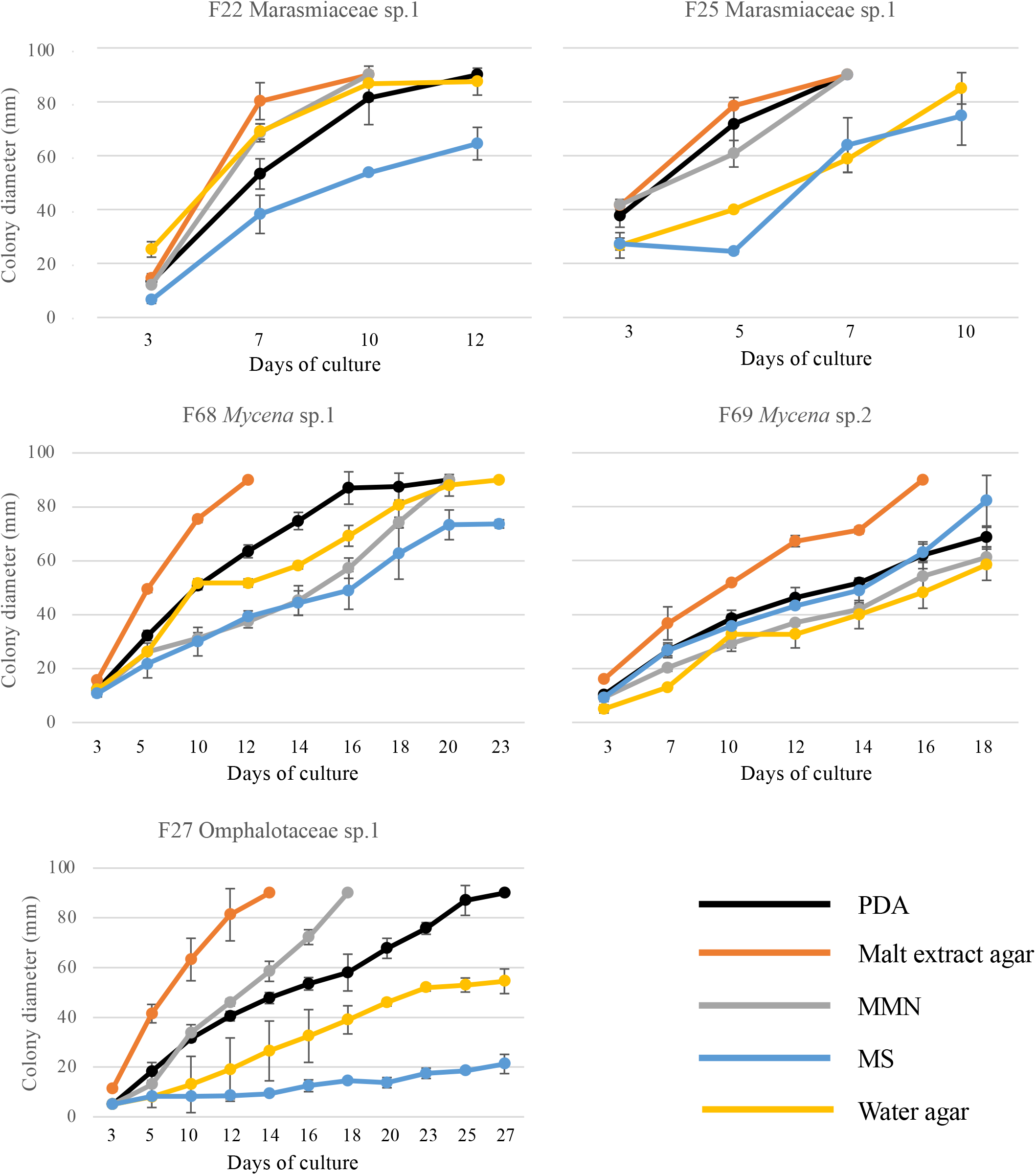
Growth of fungal isolates from *Gastrodia pubilabiata* roots on five different culture media. Mean and standard deviation are shown.

### Temperature for fungal subculture

Suitable temperature for fungal culture were evaluated by using malt extract agar medium. All fungal isolates grew fastest at 25 °C with the highest promotion in the isolate F25 (Fig. 4). There was no difference in the growth rate at 20 °C to 30 °C in the isolate F27 and F68. For all isolates, no fungal growth was found at 10 °C and 40 °C, except for the isolate F68 showing slight mycelial growth at 10 °C. All these results suggest that the culture at 25 °C would be the best for fungal subculture. Two *Mycena* strain, which induced seed germination of *Gastrodia elata*, showed the best mycelial growth at 25°C between 15°C and 30°C (Hong et al. 2002), which is consistent with the results of the present study. The appropriate growth temperature for *Mycena* depended on the climate in which the fungus is grown, with 25°C being optimal for strains of subtropical origin, and 20°C for strains of cool temperate and subalpine origin (Osono 2015). Since the fungal isolates used in this study originated from a warm temperate area, 25°C is probably appropriate.

**Fig. 4.**
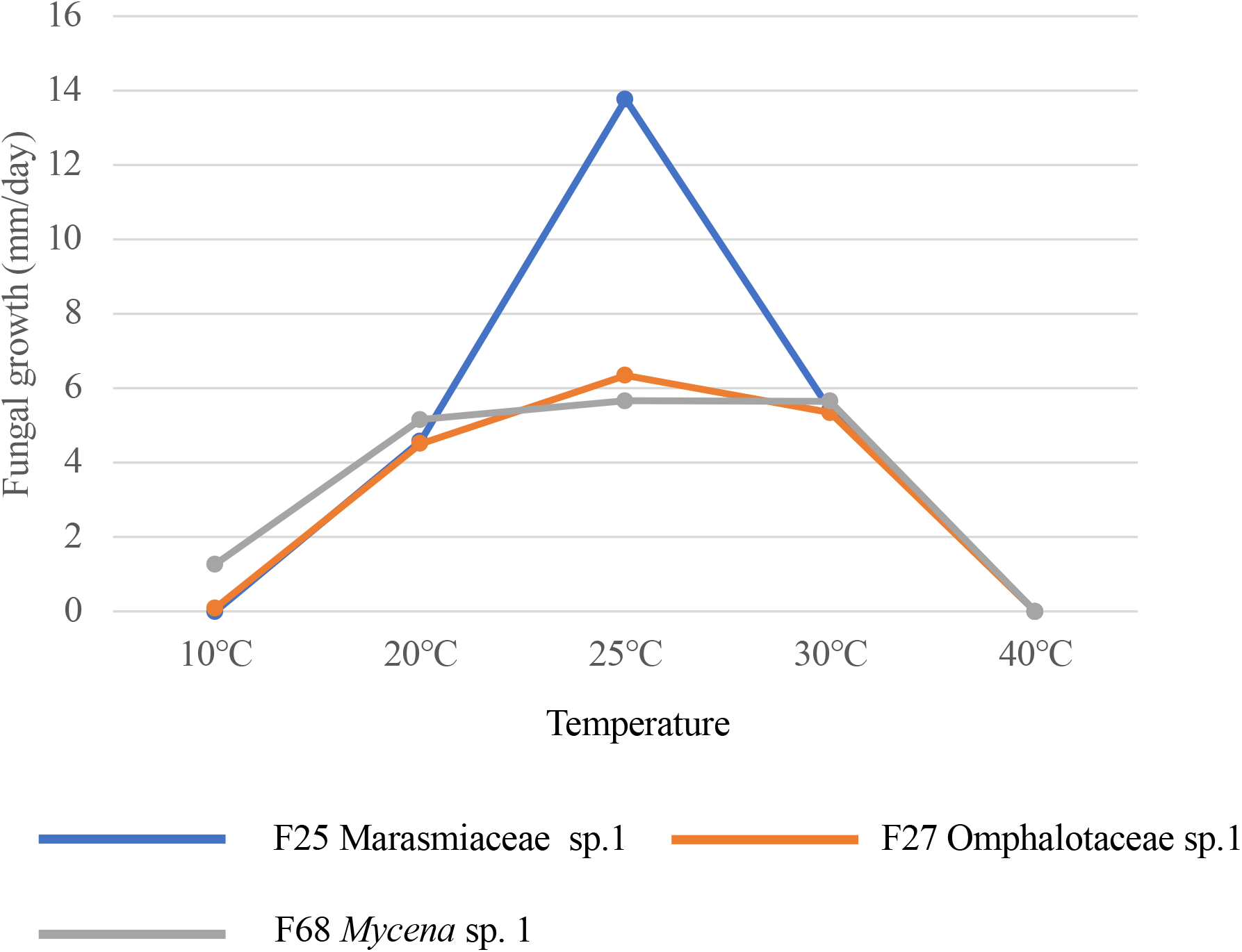
Growth of fungal isolates from *Gastrodia pubilabiata* roots under five different temperatures.

### Culture vessels for symbiotic culture

Seed germination was observed in all three vessels used in this study (Fig. 5). Seedlings were well developed in the petri dishes with leaf discs and in the glass bottles (Fig. 5A, B), and the seedlings developed to Stage 3 in the petri dishes and Stage 4 in the glass bottes. The fungal colonies grew densely in a glass bottle and three-compartment petri dish, which disturbed the observation of the seed germination process in detail. On the other hand, the seedlings in the petri dishes could be clearly distinguished on the media due to the adequate mycelial density, suggesting that this method would be suitable to observe the seed germination process and to calculate the germination rate accurately. The glass bottle would be the better method to obtain more developed seedlings within a short period.

**Fig. 5.**
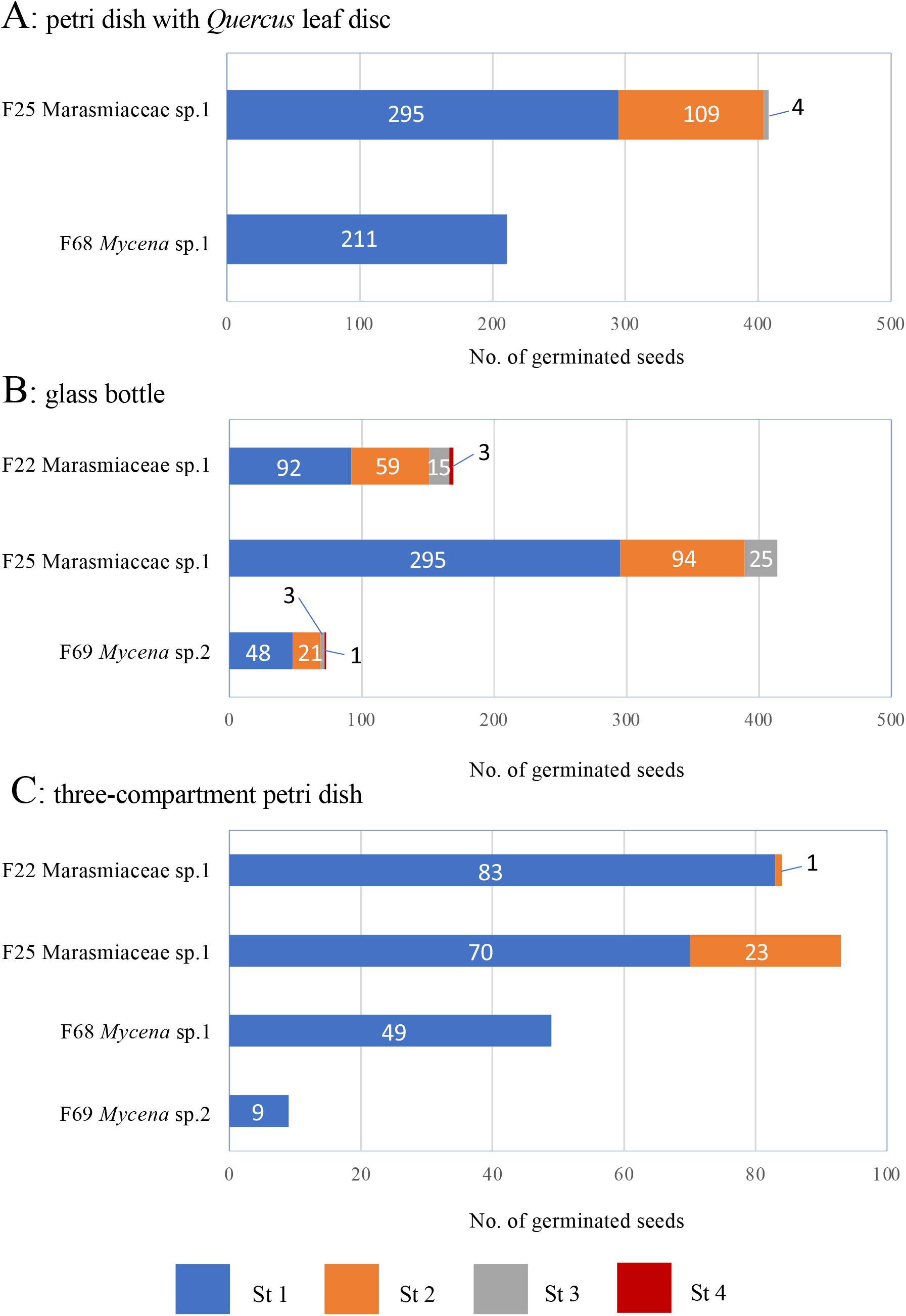
A total number of germinated seeds of *Gastrodia pubilabiata* by using three different culture vessels for symbiotic culture. Values represent a total number of germinated seeds in four replications after two months of culture.

### Organic materials for symbiotic culture

Seed germination was induced by both fungal isolates with all three organic materials, showing that these materials were suitable for symbiotic germination with *Mycena* and Marasmiaceae fungi (Fig. 6). Symbiotic seed germination of *G. elata* with *Mycena* fungi was induced on *Quercus* leaf medium, but not on bamboo leaf medium (Hong et al. 2002). The decomposition of leaf-litter by *Mycena* fungi varied with litter type (Osono 2015). Since our *Mycena* isolate F68 induced seed germination on both *Quercus* and bamboo leaves, this *Mycena* fungi may has the decomposition ability for both plant leaves.

**Fig. 6.**
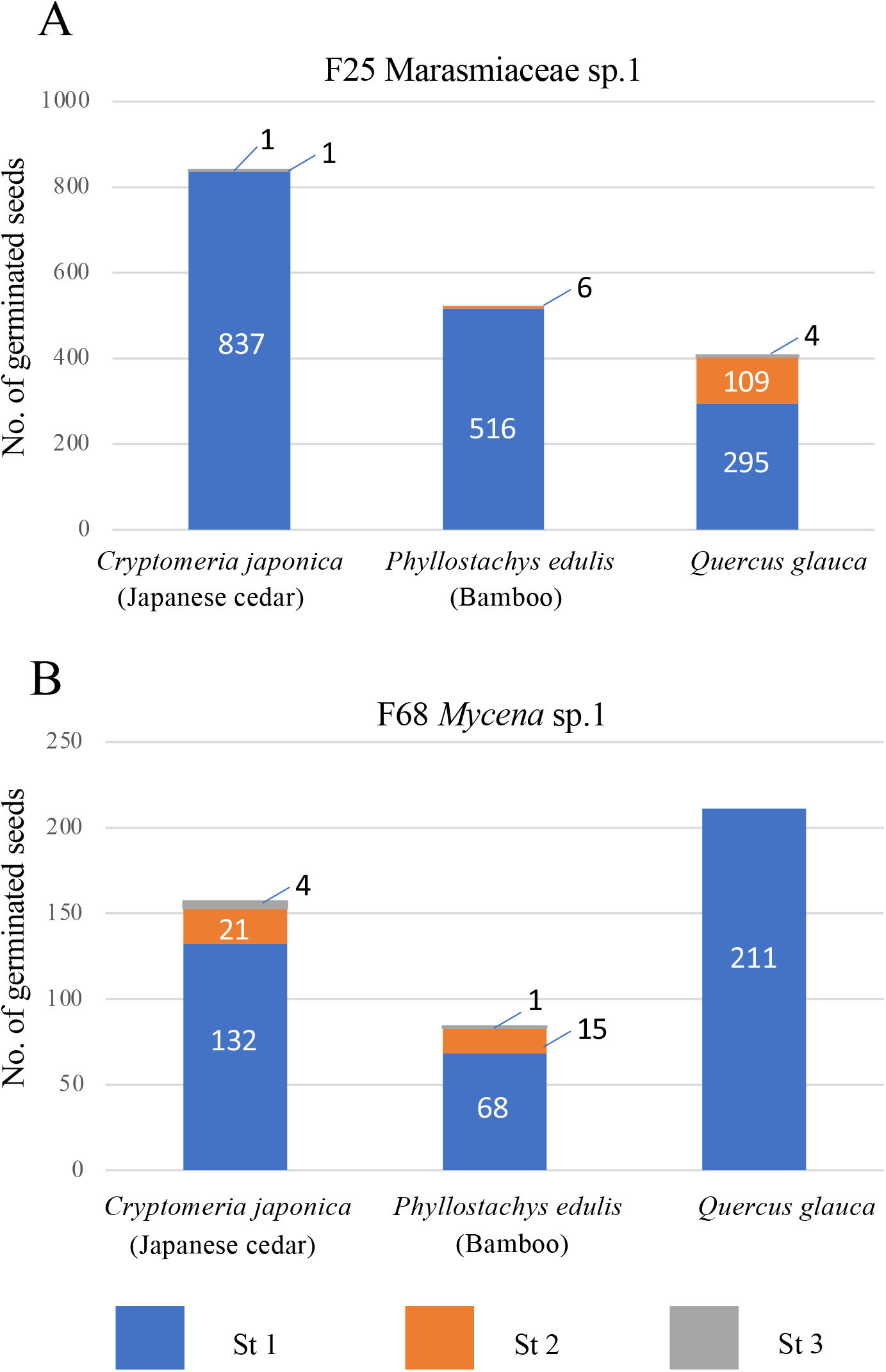
A total number of germinated seeds of *Gastrodia pubilabiata* by using three different organic materials after two months of symbiotic culture. The data for *Quercus* leaf is the same with Fig. 5.

While the seeds germinated with all three organic materials, subsequent seedling growth varied among materials. The isolate F25 with *Quercus* leaves produced the highest proportion of Stages 2 and 3, accounting for 25% of the total (Fig. 6A), while F68 with *Cryptomeria* and *Phyllostachys* leaves produced more Stages 2 and 3 seedlings than with *Quercus* (Fig. 6B). Organic material type may affect not seed germination but subsequent seedling growth. The total number of germinated seeds decreased with the growth of Stage 2 and 3 seedlings. If organic resources are concentrated on a particular individual, that individual may grow larger, but the total number of seedlings may become smaller. On the other hand, if the resource is dispersed among a large number of seedlings, the total number of seedlings may increase, but the individuals may not grow as large.

### Conclusion

This study successfully demonstrated in vitro symbiotic germination of *G. pubilabiata* with its isolates. We firstly employed Marasmiaceae and Omphalotaceae fungi for in vitro symbiotic culture of *Gastrodia* species, although various *Mycena* fungi, such as *M. osmundicola* (Xu & Guo 1989; Kim et al. 2006) and *M. anoectochila* (Guo et al. 1997), have been applied to *G. elata* culture. The results of this study indicate that not only *Mycena* but also other litter-decaying fungi can be used for in vitro culture of *Gastrodia* species. Our results also showed the optimal culture medium and temperature for subculture of Marasmiaceae, Omphalotaceae and *Mycena* isolates, in addition to suitable culture vessel and organic materials for symbiotic culture. This information will be a basis for in vitro symbiotic culture of *Gastrodia* species and also other mycoheterotrophic species that associate with leaf-litter decaying fungi. Such in vitro culture system will contribute to understanding the physiology of mycoheterotrophy. Litter-decaying fungi within Mycenaceae, Marasmiaceae and Omphalotaceae are mainly involved in mycorrhizal association of *Gastrodia* species and fungal specificities vary among plant species (Ogura-Tsujita et al. 2021). These fungal species may play an important role in seed germination, but their involvement is not clear because data on mycorrhizal association of *Gastrodia* species are mainly from adult plants. Symbiotic compatibility between *Gastrodia* species and these litter-decaying fungus can be evaluated by using in vitro symbiotic culture system, which particularly useful for identifying fungi involved in seed germination and early developmental stage. Symbiotic cultures of various litter-decaying fungi and *Gastrodia* species will reveal differences in the fungal specificity and their mechanisms.

## Acknowledgement

The authors thank to A. Kinoshita, M. Goto, K. Tanaka, N. Tanaka, Y. Yamashita and T. Yukawa for sampling and valuable advice. This study was supported by Research Grant for Plant Science from the New Technology Development Foundation.

## Notes

### Competing Interest Statement

The authors have declared no competing interest.

